# Minimized dark cataplerosis of the Calvin cycle ensures prompt photosynthetic initiation in cyanobacteria

**DOI:** 10.1101/2023.10.18.562951

**Authors:** Kenya Tanaka, Akihiko Kondo, Tomohisa Hasunuma

**Affiliations:** Engineering Biology Research Center, Kobe University, 1-1 Rokkodai, Nada, Kobe 657-8501, Japan; Graduate School of Science, Innovation and Technology, Kobe University, 1-1 Rokkodai, Nada, Kobe 657-8501, Japan; Research Center for Solar Energy Chemistry, Graduate School of Engineering Science, Osaka University, Toyonaka, Osaka 560-8531, Japan; RIKEN Center for Sustainable Resource Science, 1-7-22 Suehiro, Tsurumi, Yokohama, Kanagawa 230-0045, Japan; Department of Chemical Science and Engineering, Graduate School of Engineering, Kobe University, 1-1 Rokkodai, Nada, Kobe 657-8501, Japan

**Author notes:** Kenya Tanaka, Engineering Biology Research Center, Kobe University, 1-1 Rokkodai, Nada, Kobe 657-8501, Japan, Tomohisa Hasunuma, Engineering Biology Research Center, Kobe University, 1-1 Rokkodai, Nada, Kobe 657-8501, Japan.

## Abstract

As primary contributors to oxygenic photosynthesis, cyanobacteria intricately regulate their metabolic pathways during the diurnal cycle to ensure survival and growth. Under dark conditions, breakdown of stored energy reserves of glycogen replenishes the intermediates, especially the downstream glycolytic metabolites necessary for photosynthetic initiation upon light irradiation. The intracellular level of the intermediates is maintained throughout the dark period. However, it remains unclear how their accumulation is maintained in the dark despite the limited availability of glycogen. Here, we showed that the metabolite accumulation stability is ensured by the low activities of phosphoenolpyruvate (PEP) converting enzymes, namely PEP carboxylase and pyruvate kinase, during the dark period. Overexpression of these enzymes significantly decreased the accumulation of glycolytic intermediates after dark incubation. The oxygen evolution ability simultaneously decreased in the overexpressing strains, indicating that the dark limitation of the PEP-consuming pathway facilitates photosynthetic initiation through the maintenance of glycolytic intermediates. This finding shed light on the importance of controlling cataplerotic flux during the dark for maintaining stable operation of the Calvin cycle.

## Introduction

Cyanobacteria are oxygenic photosynthetic prokaryotes with autotrophic growth ability. In their evolutionary history, they have survived drastic environmental perturbations. Proper regulation of the dark metabolism is essential for the growth of cyanobacteria in diurnal environments (Welkie et al., 2018). During the dark period, cyanobacteria catabolize and glycogen accumulates during the light period, while the inactivation of the Calvin–Benson–Bassham (CBB) cycle minimizes energy consumption (Gurrieri et al., 2021; Diamond et al., 2015; Pittanayak et al., 2014; Saha et al., 2016). The primary role of the dark metabolism is the supply of reducing power. Carbon from glycogen catabolism is metabolized through the oxidative pentose phosphate pathway (OPPP), generating NADPH. This reducing power is crucial for avoiding oxidative stress caused by reactive oxygen species and is essential for cell survival during the night (Diamond et al., 2017).

Another role of the dark metabolism is preparation for photosynthetic initiation. A wild-type strain was able to maintain the accumulation of the CBB cycle intermediates throughout the dark period (Diamond et al., 2017). The inability of glycogen breakdown impairs photosynthetic activation after dark incubation due to the depletion of CBB cycle intermediates (Shimakawa et al., 2014; Shinde et al., 2020). While the OPPP is the main pathway generating glycogen intermediates (Yang et al., 2002; Wan et al., 2017), other glycolytic shunts such as the Entner–Doudoroff (ED) pathway have been suggested to help compensate for their depletion (Makowka et al., 2020). It has been proposed that the CBB cycle is not completely autonomous, but that it rather relies on anaplerotic reactions (OPPP and other glycolytic shunts) similar to the other cyclic pathways such as the tricarboxylic acid (TCA) cycle (Makowka et al., 2020). However, it remains unclear how the supplied CBB cycle intermediates are maintained even though glycogen availability is limited during the dark period.

In our recent study, we showed that the dark accumulation of glycolytic intermediates, specifically three phosphorylated acid intermediates (3-phosphoglycerate; 3PG, 2-phosphoglycerate; 2PG, and phosphoenolpyruvate; PEP), facilitates the efficient initiation of photosynthesis (Tanaka et al., 2023). Anaplerotic reactions for the CBB cycle, such as glycogen catabolism and the OPPP, have been suggested to replenish these three glycolytic acid intermediates. However, the impact of cataplerotic reactions of the three intermediates on the accumulation of the three acid intermediates is unknown. We hypothesized that the limitation in the dark consumption of PEP, the downstream metabolite among the three intermediates, leads to the accumulation of glycolytic intermediates, enabling rapid start of photosynthesis. In cyanobacteria, pyruvate kinase (Pyk) and PEP carboxylase (PepC) are the main metabolic enzymes converting PEP. The present study investigated the effects of Pyk or PepC overexpression on the accumulation of metabolites as well as photosynthetic rates using the cyanobacterium *Synechocystis* sp. PCC 6803. Our findings revealed that an elevated PEP conversion rate during the dark period leads to an inappropriate decrease in the concentration of glycolytic intermediates, impairing the smooth initiation of photosynthesis during the transition to the light period.

## Results

In our previous study, we found that overexpression of the gene encoding PepC (*ppc*) increased the fermentation production of succinate under dark anaerobic conditions (Hasunuma et al., 2016). In the present study, to investigate the impact of *ppc* overexpression on dark metabolism, we measured the intermediate levels in the central metabolic pathway during dark adaptation in the presence of 50 mM NaHCO_3_, a substrate for PepC (Figure 1, Supplemental Figure S1). As a result, the glycolytic acid intermediates were significantly decreased in the Ppc-ox strain compared to those in the wild type (WT). As glycolytic acid intermediates are the main substrates for photosynthetic initiation immediately after the dark-to-light transition (Tanaka et al., 2023), their depletion in the Ppc-ox strain can decrease the oxygen evolution rate immediately after light irradiation. In the Ppc-ox strain, the cells adapted to darkness for 68 hours exhibited a significantly reduced oxygen evolution rate compared to those of the WT (Figure 2). On the other hand, the effective quantum yield of PSII [Y(II)] in the Ppc-ox strain was consistently lower than that in the WT throughout the experiment (Figure 2). Y(II) is known to increase through the alternative electron flow mediated by flavodiiron proteins in cyanobacteria (Shimakawa et al., 2015). The alternative electron flow may be suppressed in the Ppc-ox strain for an unknown reason. Our results indicated that in the Ppc-ox strain, the enhanced PEP consumption by PepC decreases the accumulation of glycolytic acid intermediates, resulting in insufficient substrate for CO_2_ fixation and a decrease in oxygen evolution rate.

**Figure 1.**
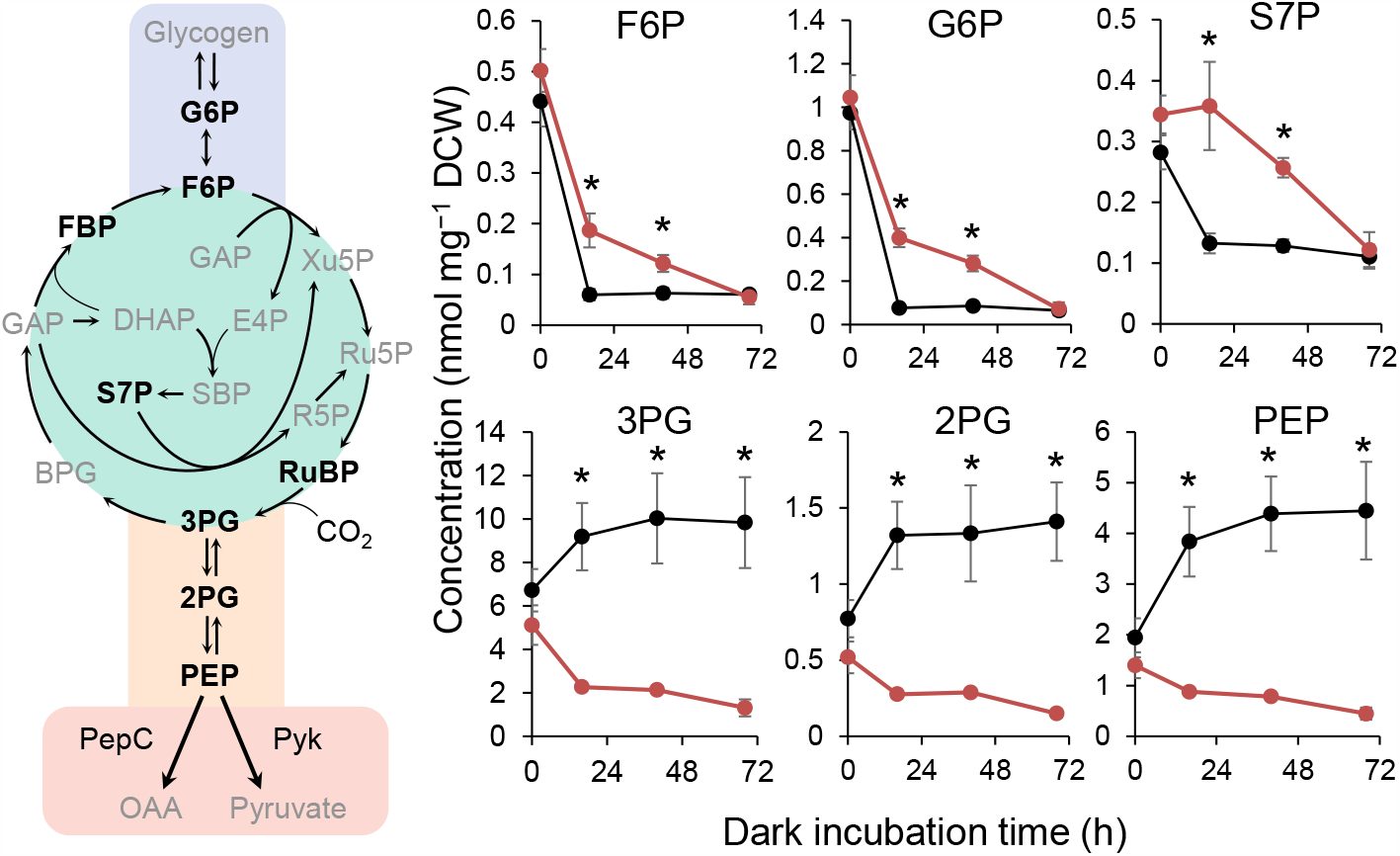
Effect of phosphoenolpyruvate carboxylase (PepC) overexpression on the intracellular metabolite concentration during dark incubation (black: wild type, red: PepC-ox strain). Metabolic pathways shown on the left are classified by background color (blue: glycogen pathway, green: Calvin-Benson-Bassham cycle, orange: glycolysis, red: phosphoenolpyruvate consuming pathways). Metabolites written in bold letters are the measured intermediates. The other metabolites were peak-unidentified (BPG, SBP, and OAA), unmeasured (glycogen), or under the detection limit (others < 0.01 nmol mg−1 DCW). Values are shown as mean ± SD (bars) of three biological replicates. Significant differences between the strains were evaluated by a two-tailed non-paired Student’s t test (* *p* < 0.05). Abbreviations: BPG; 1,3-bisphosphoglycerate; DCW, dry cell weight; DHAP, dihydroxyacetone phosphate; E4P, erythrose 4-phosphate; FBP, fructose 1,6-bisphosphate; F6P, fructose 6-phosphate; GAP, glyceraldehyde 3-phosphate; G6P, glucose 6-phosphate; PEP, phosphoenolpyruvate; Pyk, pyruvate kinase; 2PG, 2-phosphoglycerate; 3PG, 3-phosphoglycerate; RuBP, ribulose 1,5-bisphosphate; Ru5P, ribulose 5-phosphate; R5P, ribose 5-phosphate; SBP, sedoheptulose 1,7-bisphosphate; S7P, sedoheptulose 7-phosphate; Xu5P, xylulose 5-phosphate.

**Figure 2.**
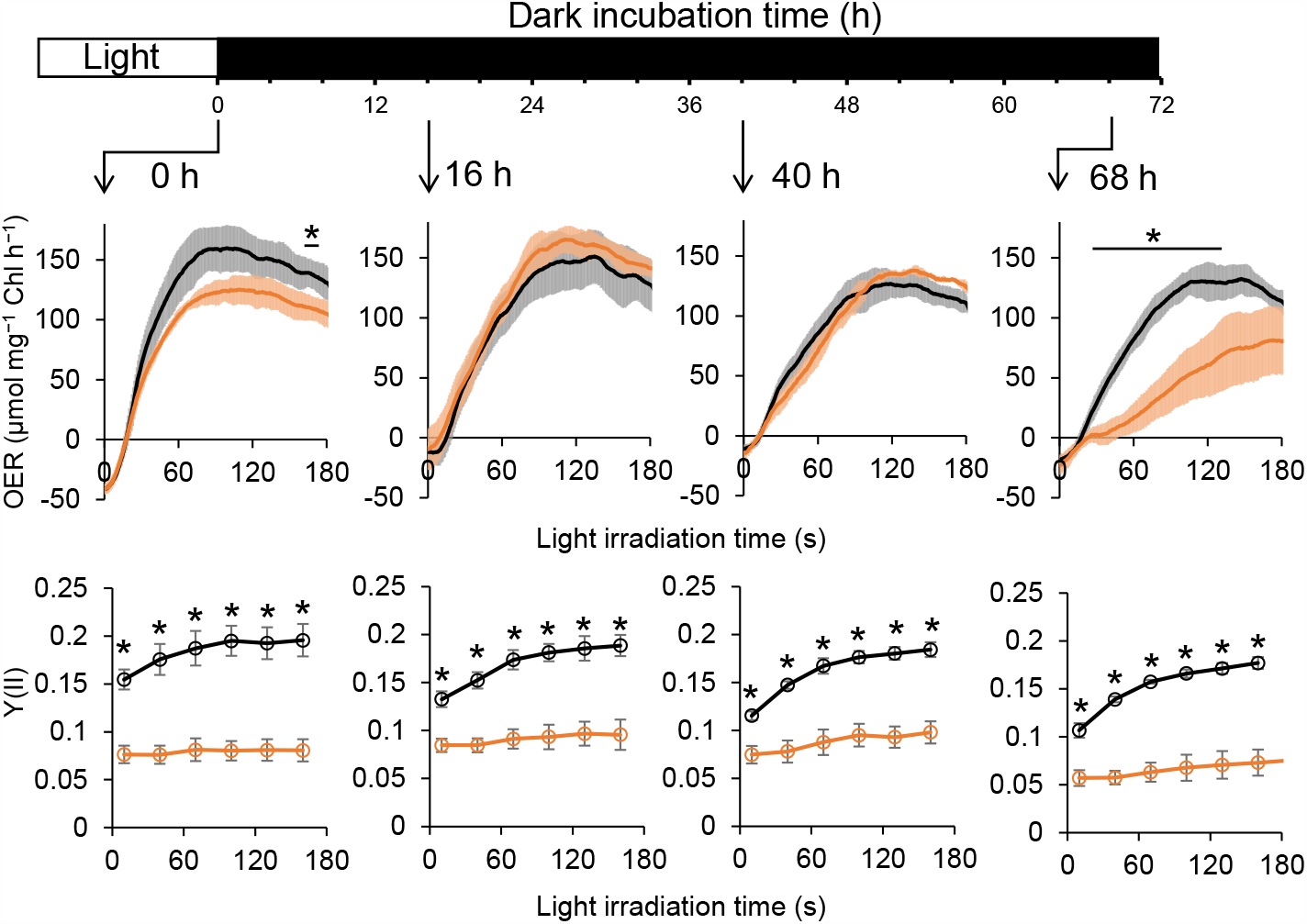
Photosynthetic initiation after dark incubation in the wild type (black) and Ppc-ox strain (orange). Dark-incubated cells were illuminated for measuring the oxygen evolution rate (OER) and effective quantum yield of PSII [Y(II)]. Values are shown as mean ± SD (bars) of three biological replicates. Significant differences between the strains were evaluated by a two-tailed non-paired Student’s t test (* *p* < 0.05).

The conversion of PEP to pyruvate is another PEP consumption pathway, and Pyk has been suggested to be the enzyme controlling the flux of this pathway (Jazmin et al., 2017; Nishiguchi et al., 2019). To examine whether the low Pyk activity contributes to the maintenance of central intermediates and initiation of photosynthesis, we investigated the influence of Pyk overexpression. The endogenous Pyk gene (*sll0587*) was introduced under the *trc* promoter into *Synechocystis* by homologous recombination. Gene insertion was confirmed by PCR (Supplemental Figure S2). Pyk activity was over 20 times higher in the Pyk-ox strain than in the WT under both light and dark conditions, confirming the overexpression of Pyk; however, the diurnal changes in Pyk activity were small and not significant (Figure 3A). Furthermore, we investigated the effect of Pyk overexpression on metabolism (Figure 3B, Supplemental Figure S3). In WT, the accumulation of glycolytic acid intermediates was higher during the dark period than during the light period, suggesting their active accumulation for photosynthetic start. In contrast, in Pyk-ox, the accumulation of glycolytic acid intermediates during the dark period was significantly lower than that in WT (Figure 3B). Glycogen accumulation was significantly lower in Pyk-ox than in WT under dark conditions (Figure 3C). This suggested that Pyk overexpression promoted glycogen catabolism and implied that Pyk reaction is one of the bottlenecks in the downstream metabolic pathways. Additionally, in Pyk-ox, the adenylate energy charge was significantly lower than that of WT during the dark period (Figure 3D). Pyk reaction catalyzes the conversion from ADP to ATP. ATP is consumed and converted into AMP when pyruvate is converted back to PEP by PEP synthase. The series of Pyk and PEP synthase reactions forms a fertile cycle where ADP is converted to AMP, potentially leading to an increase in AMP. However, the reason for the decrease in ATP instead of a decrease in ADP is not clear.

**Figure 3.**
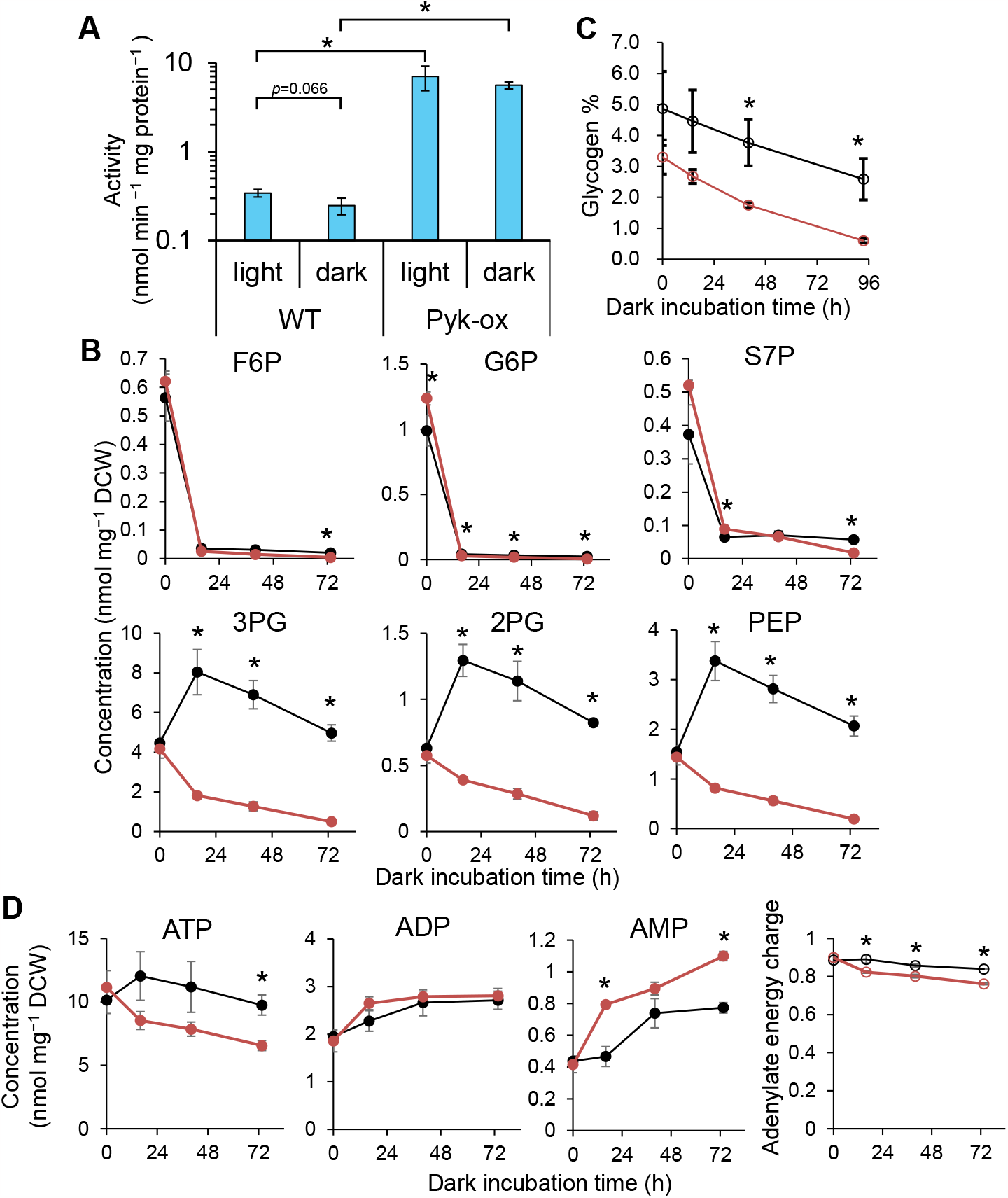
Effect of pyruvate kinase gene overexpression. (A) Pyruvate activity in the cell crude extract of the wild type (WT) and Pyk-ox strain. (B) Intracellular metabolite concentration during dark incubation. (C) Glycogen content. (D) Adenosine phosphate content and adenylate energy charge (black: WT, red: Pyk-ox strain). Values are shown as mean ± SD (bars) of three biological replicates. Significant differences between the strains were evaluated by a two-tailed non-paired Student’s t test (* *p* < 0.05).

Subsequently, we measured oxygen evolution to examine the impact of metabolic changes caused by Pyk overexpression on photosynthetic initiation. While the Pyk-ox strain adapted to light conditions (t=0) showed similar oxygen evolution activity to that of WT, with increasing dark adaptation time, the oxygen evolution rate of the Pyk-ox strain significantly decreased compared to that of WT (Figure 4). Moreover, the Y(II) values of 74 h dark-adapted cells were significantly lower in the Pyk-ox strain than in the WT (Figure 4). These results indicated that the cataplerotic flux enhanced by Pyk overexpression depletes the glycolytic acid intermediates during the dark period, leading to a decrease in the oxygen evolution rate immediately after light exposure. The decrease in Y(II) was also suggested to be caused by the deficiency of glycolytic acid intermediates. However, the relatively minor impact compared to the decrease in oxygen evolution rate may be attributed to the operation of alternative electron flow.

**Figure 4.**
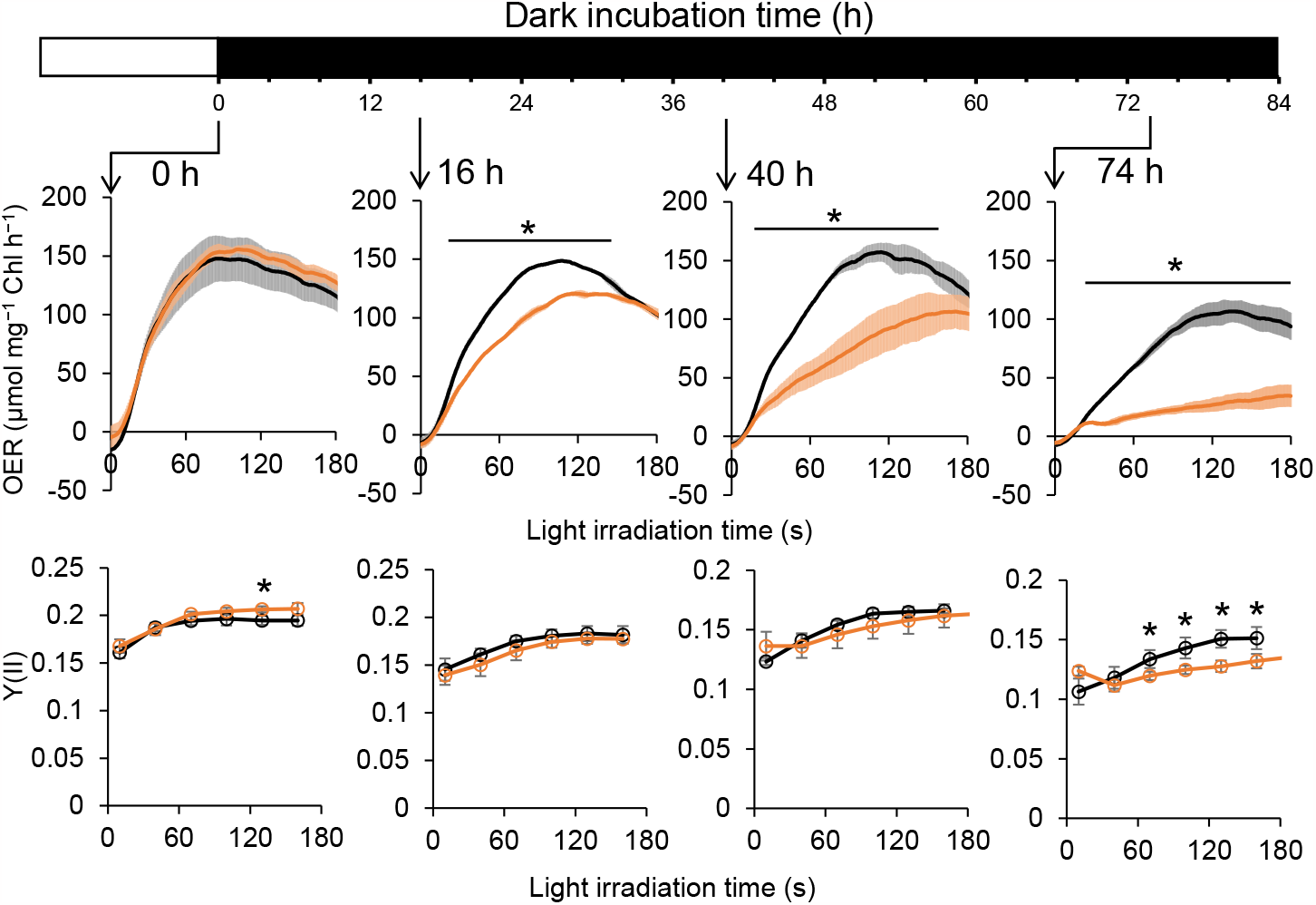
Photosynthetic initiation after dark incubation in the wild type (black) and Pyk-ox strain (orange). Values are shown as mean ± SD (bars) of three biological replicates. Significant differences between the strains were evaluated by a two-tailed non-paired Student’s t test (* *p* < 0.05).

Finally, based on the obtained data, we examined the relationship between the 3PG accumulation and the oxygen evolution rate 90 seconds after light exposure. As shown in Figure 5, it became evident that the oxygen evolution rate significantly decreased below a certain threshold of 3PG concentration. Thus, the accumulation of glycolytic acid intermediates is crucial for photosynthetic activity, and the maintenance of acid intermediate accumulation is dependent on the low activities of Pyk and PepC.

**Figure 5.**
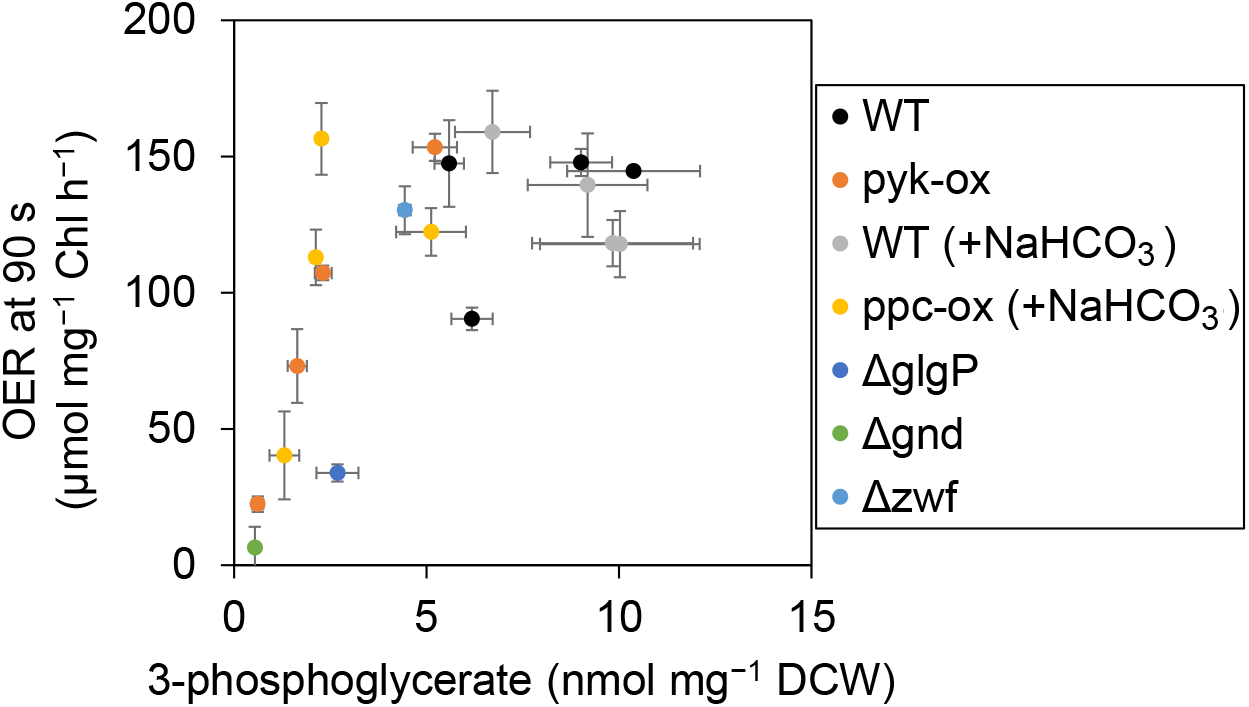
Relationship between intracellular 3-phosphoglycerate (3PG) concentration and oxygen evolution rate (OER) at 90 seconds after light irradiation. The data of ΔglgP, ΔgndP, and Δzwf strains from Tanaka et al., (2023) are shown.

## Discussion

While anaplerotic reactions for the CBB cycle have been demonstrated to be crucial for maintaining the reducing power and supplying CBB cycle intermediates in the dark metabolism, little is known about the impact of cataplerotic reactions. This study revealed that low activities of Pyk and PepC reactions result in the accumulation of three glycolytic acid intermediates during dark period, and these intermediates are essential for the rapid activation of the CBB cycle (Figure 6). Pyk activity is likely to remain low throughout the light-dark cycles (Figure 3A), while the expression of the *ppc* gene is controlled under the diurnal cycle, peaks during the light period, and decreases during the dark period in *Synechocystis* (Saha et al., 2016). Among the pathways consuming PEP, PepC has the largest flux during the light period (Young et al., 2011; Jazmin et al., 2017). Downregulation of PepC during the dark period along with maintaining low Pyk is likely to help maintain the optimal metabolic state during the dark period. The control of these cataplerotic reactions may facilitate metabolic switching from dark to light conditions in photosynthetic organisms.

**Figure 6.**
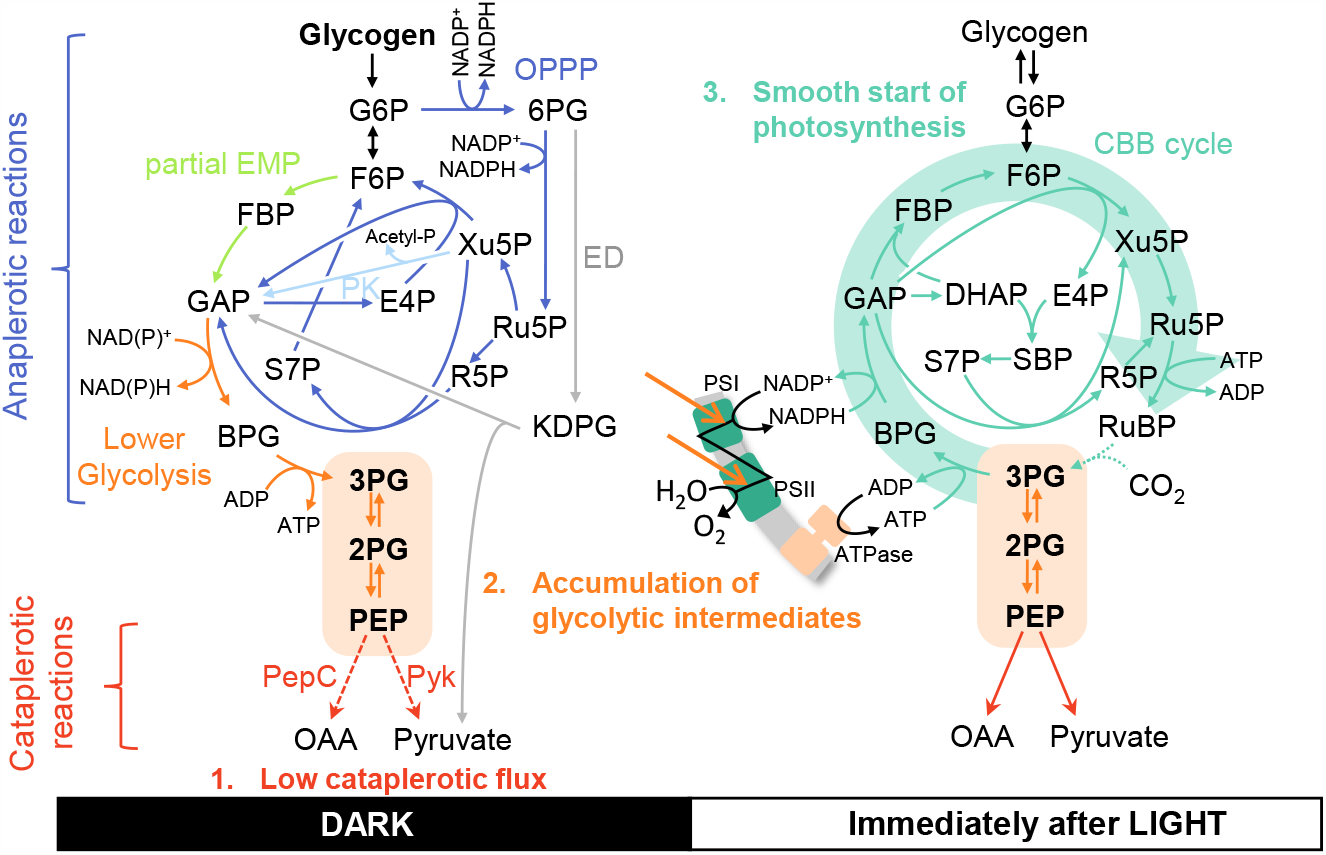
Proposed mechanism of dark accumulation of glycolytic intermediates and rapid initiation of the CBB cycle (green arrows). During the dark period, glycolytic intermediates (3PG, 2PG, and PEP) are replenished from glycogen through OPPP (blue arrows) and lower glycolysis (orange arrows). The ED (gray arrows), PK (light blue arrows), and partial EMP (light green arrows) pathways may contribute to the replenishment as well. The thick green arrow in the right panel indicates the metabolic flow during photosynthetic initiation, and the dashed green arrow indicates the CO_2_ fixation flux because activation of CO_2_ fixating enzyme Rubisco is delayed after light irradiation. Abbreviations: CBB, Calvin–Benson–Bassham; ED, Entner–Doudoroff; EMP, Embden–Meyerhof–Parnas; KDPG, 2-keto-3-deoxy-6-phosphogluconate; OPPP, oxidative pentose phosphate pathway; PK, phosphoketolase; 6PG, 6-phosphogluconate. The other abbreviations are the same as in Figure 1.

In the WT, the transition from light to dark led to a decrease in glycogen and sugar phosphates such as glucose-6-phosphate and an increase in downstream glycolytic intermediates, namely 3PG, 2PG, and PEP (Figures 1 and 3). This implied that glycogen and sugar phosphates are converted to the downstream glycolytic intermediates under dark conditions through metabolic pathways discussed below (Figure 6). OPPP is an essential pathway supplying the necessary reducing power and metabolic intermediates from glycogen to sustain the cellular functions during the dark (Diamond et al., 2017; Wan et al., 2017). The contribution of the partial Embden– Meyerhof–Parnas (EMP) pathway (fructose 6-phosphate to glyceraldehyde-3-phosphate) during the dark should be minor. Although an early study on the metabolic flux in *Synechocystis* showed that the partial EMP is active under dark heterotrophic conditions (Yang et al., 2002), another study observed no partial EMP flux under similar conditions (Wan et al., 2017). Moreover, the deletion of phosphofructokinase, the key enzyme of the EMP pathway, was reported to not affect the reactivation of the CBB cycle (Makowka et al., 2020). While the deletion of the ED pathway delayed the onset of the CBB cycle after the short dark pulse (Makowka et al., 2020), it does not have a significant metabolic flux under dark heterotrophic conditions (Wan et al., 2017). The ED pathway may be active during the onset of darkness. Phosphoketolase (PK) converts sugar phosphates to glyceraldehyde-3-phosphate (GAP) and acetyl phosphate. In dark heterotrophic conditions, PK is necessary for acetate production from exogenous sugars (Xiong et al., 2015). Recently, it has been revealed that the consumption of sugar phosphates by PK immediately after the light-to-dark transition halts the CBB cycle (Lu et al., 2023). The GAP generated by PK potentially contributes to the accumulation of glycolytic acid intermediates. All the replenishment pathways mentioned above (OPPP, partial EMP, ED, and PK pathways) generate GAP from sugar phosphates (Figure 6). GAP can be converted to 3PG by glyceraldehyde-3-phosphate dehydrogenase (GAPDH) and phosphoglycerate kinase (PGK). *Synechocystis* harbors two GAPDH genes, namely *gap1* and *gap2*, which differ in cofactor specificity toward NAD and NADP (Koksharova et al., 1998). Furthermore, the regeneration of GAP through the reduction in 3PG is inactivated by CP12 (Tamoi et al., 2005; McFarlane et al., 2019). Taken together, the metabolic system of cyanobacteria appears to be designed in a way that downstream glycolytic metabolites accumulate easily in dark conditions.

In photosynthetic bio-production by cyanobacteria utilizing light energy and CO_2_, the PEP conversion pathway is a target for metabolic engineering to increase the production of target compounds. Overexpression of *ppc* enhances succinate production, while *pyk* overexpression increases the production of 2,3-butanediol (23BD) and isobutyraldehyde (IBA) (Hasunuma et al., 2016; Oliver and Atsumi, 2015; Jazmin et al., 2017). These observations suggested that the PEP consumption pathways are recognized as bottleneck reactions in the production of downstream metabolites. Among these studies, Oliver and Atsumi (2015) demonstrated that enhancing the cataplerotic flux through overexpression of *pgm, eno*, and *pyk* genes in a 23BD-producing strain led to a decrease in total carbon productivity (sum of growth and 23BD productivity) in *Synechococcus elongatus* PCC 7942. The authors proposed that this decrease might be related to an excessive carbon flux toward pyruvate, potentially depleting the CBB cycle intermediates, although the concentrations of these intermediates were not measured. On the contrary, a single overexpression of *pyk* was found to increase the total carbon productivity. This photosynthetic enhancement is considered to be a phenomenon that occurs when expressing and increasing the sink metabolism; however, the precise mechanisms remain unknown (Santos-Merino et al., 2021).

In the present study, under continuous light and light-dark conditions, the growth rates of the WT and Pyk-ox were similar (Supplemental Figure S4). While a reduction in photosynthetic activity during photosynthetic activation and a decrease in adenylate energy charge during the dark period were observed in the Pyk-ox strain (Figures 3D and 4), its impact on growth appears to be limited. A further increase in the cataplerotic flux of the CBB cycle is expected to affect the growth and photosynthetic efficiency. As the industrial application of photosynthetic organisms will require efficient growth and production in outdoor conditions with light-dark environments, further investigation is required to elucidate the relationship between dark metabolism and Calvin cycle operation.

## Materials and Methods Cyanobacteria strains

All strains were constructed in a glucose-tolerant (GT) strain *Synechocystis* sp. PCC 6803. The Ppc-ox and vector control strains (denoted as WT with kanamycin resistant cassette) constructed in Hasunuma et al., (2016) were used for the experiments. For the construction of a Pyk overexpression strain, the endogenous *pyk* gene sll0587 was amplified from *Synechocystis* genomic DNA by PCR using the primer set 5’-GGAAACAGACCCATATGCCAGCCCTGATTAACCC-3’/5’-TAACCTGCAGGTCGACGCAACTATGGATTTGAGGCATAGG-3’. The resulting fragment was integrated into *NdeI*/*SalI* digested pSKtrc-slr0168 constructed in Hasunuma et al., (2016) using an In-Fusion HD Cloning Kit (TakaraBio, Shiga, Japan) to yield pSKtrc-slr0168/sll0587. The GT strain was transformed with pSKtrc-slr0168/sll0587 to yield the strain Pyk-ox. The chromosomal integration of pyk was confirmed by PCR using the primer set 5’-ATGGCACCGATGCGGAATCCCAACAGATTGCCTTTGAC-3’/5’-CACGTTGGGTCCCAAGTTTGTGCTGTGGCTGATGCCAT-3’.

### Culture conditions

All strains were grown in the BG-11 liquid medium. The cells were pre-cultured for five days in 200-mL flasks containing 50 mL of the BG-11 medium and 20 mM of HEPES-NaOH (pH 7.5) in Bioshaker BR-43FL (TAITEC, Saitama, Japan) under continuous light irradiation (LC-450EXP; TAITEC) at the light intensity of 30 μmol m^-2^ s^-1^ with 100 rpm agitation at 30 °C. For the main culture, the pre-cultured cells were inoculated into the fresh 70 mL BG-11 medium containing 20 mM HEPES-NaOH (pH 7.5) to the initial optical density of 0.05 at 730 nm (OD_730_). Cultivation was performed in the same way as pre-culture. The cell density was monitored by OD_730_. Dry cell weight (DCW) was determined by a linear correlation with OD_730_ values.

### Metabolite analysis

Metabolite sampling was started with a six-day-old main culture (equivalent to OD_730_: 2–3). The cells were filtered from the calculated volume of the culture (10/OD_730_ in mL) using 1 μm pore size polytetrafluoroethylene membrane (Millipore, Billerica, MA). Intracellular metabolites were extracted and analyzed by capillary electrophoresis-mass spectrometry (Agilent G7100; MS, Agilent G6224AA LC/MSD TOF; Agilent Technologies, Palo Alto, CA) as described previously (Hasunuma et al., 2016).

### Glycogen analysis

The cells were harvested by centrifugation at 8,000 ×g for 5 min from the calculated volume of the culture (15/OD_730_ in mL). The cell pellet was lyophilized by a freeze dryer (FDM-1000; EYELA, Tokyo, Japan). Glycogen extraction and analysis were performed as described previously (Hasunuma et al., 2016).

### Measurement of oxygen evolution and chlorophyll fluorescence

The cells were harvested by centrifugation at 8,000 ×g for 5 min from the calculated volume of the culture (10/OD_730_ in mL), followed by resuspension into the fresh 2-mL BG-11 medium. Oxygen evolution was measured using a 1-mL cell suspension with actinic light at the light intensity of 200 μmol m^-2^ s^-1^ irradiated by PAM-2500 (Waltz, Effeltrich, Germany) in a Clark-type oxygen electrode (Hansatech Instruments, Norfolk, UK). Chlorophyll fluorescence was simultaneously detected by PAM-2500. The apparent oxygen evolution rate was normalized using the chlorophyll *a* concentration of the cell suspension. The concentration of chlorophyll *a* was determined according to Shimakawa et al., (2014).

### Measurement of pyruvate kinase activity

The cells were harvested by centrifugation at 8,000 ×g for 5 min from the calculated volume of the culture (10/OD_730_ in mL), followed by resuspension into 200 μL of phosphate-buffered saline (PBS). The cells were disrupted by a multi-beads shocker (Yasui Kikai Co., Osaka, Japan) at 2700 rpm with ON 60 sec and OFF 120 sec for eight cycles. The extract was centrifuged at 10,000 ×g for 10 min. The supernatant was applied for the subsequent enzymatic activity assay. Pyk activity was measured using a pyruvate kinase activity assay kit (MAK072-1KT; Merck, Darmstadt, Germany) according to the manufacturer’s instructions. The reaction was monitored through fluorescence (excitation: 535 nm, emission: 587 nm) every 1 min using Infinite 200 PRO (TECAN, Männedorf, Switzerland).

## Acknowledgements

We would like to thank Editage (www.editage.jp) for English language editing.

## Author Contributions

K.T. designed the study. K.T. performed the experiments, analyzed the data and wrote the manuscript. T.H and A.K supervised the project, and contributed to interpretation of the data and edit the manuscript.

